# Vascularized tumor organoids enable immunotherapy testing in tumor-remodeled stroma

**DOI:** 10.64898/2026.01.23.701300

**Authors:** Reiner Wimmer, Rok Krese, Louisa Henniger, Steve Niu, Nadege Lagarde, Timon Keller, Inga Clausen, Fabian Köchel, Claudia Ferrara Koller, Inja Waldhauer, Marina Bacac, Nikolche Gjorevski, Fjodor Melnikov, Remi Villenave

## Abstract

Accurately predicting whether a new molecule will be safe and effective in humans remains a fundamental challenge in biomedical research, requiring models that can reliably inform clinical outcomes. In oncology, drug attrition rates remain disproportionately high compared to other therapeutic areas, largely due to the poor translatability of conventional preclinical models that fail to capture the complexity of human immune-tumor-stroma interactions (1, 2). Here, we describe human vascularized organoids (VascO), formed through self-assembly of an immune cell perfused capillary network and stromal compartment, interfacing with patient-derived colorectal cancer organoids that enable the combined safety and efficacy assessment of tumor-targeted immunotherapies. VascO generates functional and integrated vascular networks within tumor organoids that sustain organoid growth and recapitulate the morphology and functionality of tumor vasculature. Single-cell transcriptomics reveals that the tumor organoids-educated VascO niche reprograms otherwise healthy fibroblasts into canonical cancer-associated fibroblast states, adopting myofibroblastic or inflammatory signatures, and faithfully mirroring donor-specific tumor cues. In parallel, VascO tumor vessels display an expanded tip-cell program, aberrant morphology, and VEGF/ANG2-driven permeability, recapitulating defining hallmarks of tumor vasculature. Immune perfusion with a tumor-targeted T cell bispecific antibody induces dynamic immune trafficking, infiltration and engagement of effector cells with the tumor tissue, leading to its effective killing. A protease-activated variant restricts T cell activation to the tumor microenvironment, retaining potent tumoricidal activity while sparing donor-matched healthy colon organoids, and thereby improving the therapeutic index. VascO thus models a patient-specific, tumor-educated microenvironment that is amenable to assessing the combined safety and efficacy of new drug candidates and provides a scalable platform to improve the preclinical assessment of innovative cancer immunotherapies.

## Main

Generally recognized as a primary challenge in cancer immunotherapy the establishment of pre-clinical models that facilitate the development and testing of cancer immunotherapies by accurately predicting human clinical response has proven elusive. While conventional animal models, particularly rodents, have been successfully applied to assess the efficacy and safety of cancer immunotherapies, their ability to faithfully recapitulate human cancer biology and patient response to treatment is regularly called into question (3–6). Moreover, the increasing complexity of novel cancer immunotherapy mechanisms of action (7–9), expanding therapeutic modalities (10), and human-specific targets and pathways (11) further challenge their translational value. Cancer cell lines, another workhorse of cancer research owing to their expression of human targets and compatibility with high-throughput studies, have similarly played a central role in oncology drug discovery. However, as reductionist tumor models, they are fundamentally limited in their ability to recapitulate human tissue responses and predict clinical outcomes (12, 13). In parallel, momentum towards more predictive, human-relevant systems in drug development is being amplified by recent global regulatory calls to reduce and eventually phase out animal models for preclinical assessment of monoclonal antibodies, lending new urgency to a longstanding challenge (14, 15).

To address this translational gap, patients-derived tumor organoids have emerged as attractive preclinical cancer models due to their ability to maintain histological, genetic and transcriptomics features of original tumor tissues (16), reflect inter and intra-tumor heterogeneity (17, 18) and display comparable sensitivity to anti-tumor therapies as their parental tumor tissues (19–22). However, to enable the testing of cancer immunotherapies, the presence of an effector immune population and a stromal compartment comprised of fibroblasts, functional vasculature and a 3D extracellular matrix scaffold that support the spatial organization of an in vitro tumor microenvironment and mediate immune tumor infiltration is essential. Because vascularizing organoids by animals engrafting is impractical for drug testing applications, pioneering methods have been developed that generate self-assembled, perfusable vascular network with stromal support under scalable, controllable conditions (23–25), with limited proof of concept studies applying these methods to healthy organoids (26, 27). While these methods offer to functionally augment organoids with stromal interaction and perfusable microvasculature, thereby allowing immune cells circulation, their extension to patient-derived tumor organoids with functional perfusion and application to therapeutic testing has yet to be demonstrated (28, 29).

Thus, we created a functional human vascularized organoid (VascO) platform in which a self-assembled, perfusable vascular network integrates with patient-derived tumour organoids, enabling controlled delivery, trafficking, and tissue infiltration of human immune cells (Fig. 1a). We investigated relative cell impact on each compartment using single cell sequencing and used VascO to recapitulate pharmacologically driven T cell tumor infiltration and killing, demonstrating its applicability for the combined safety and efficacy assessment of two successive generations of T cell bispecific antibodies to help estimate a therapeutic window.

**Figure 1.**
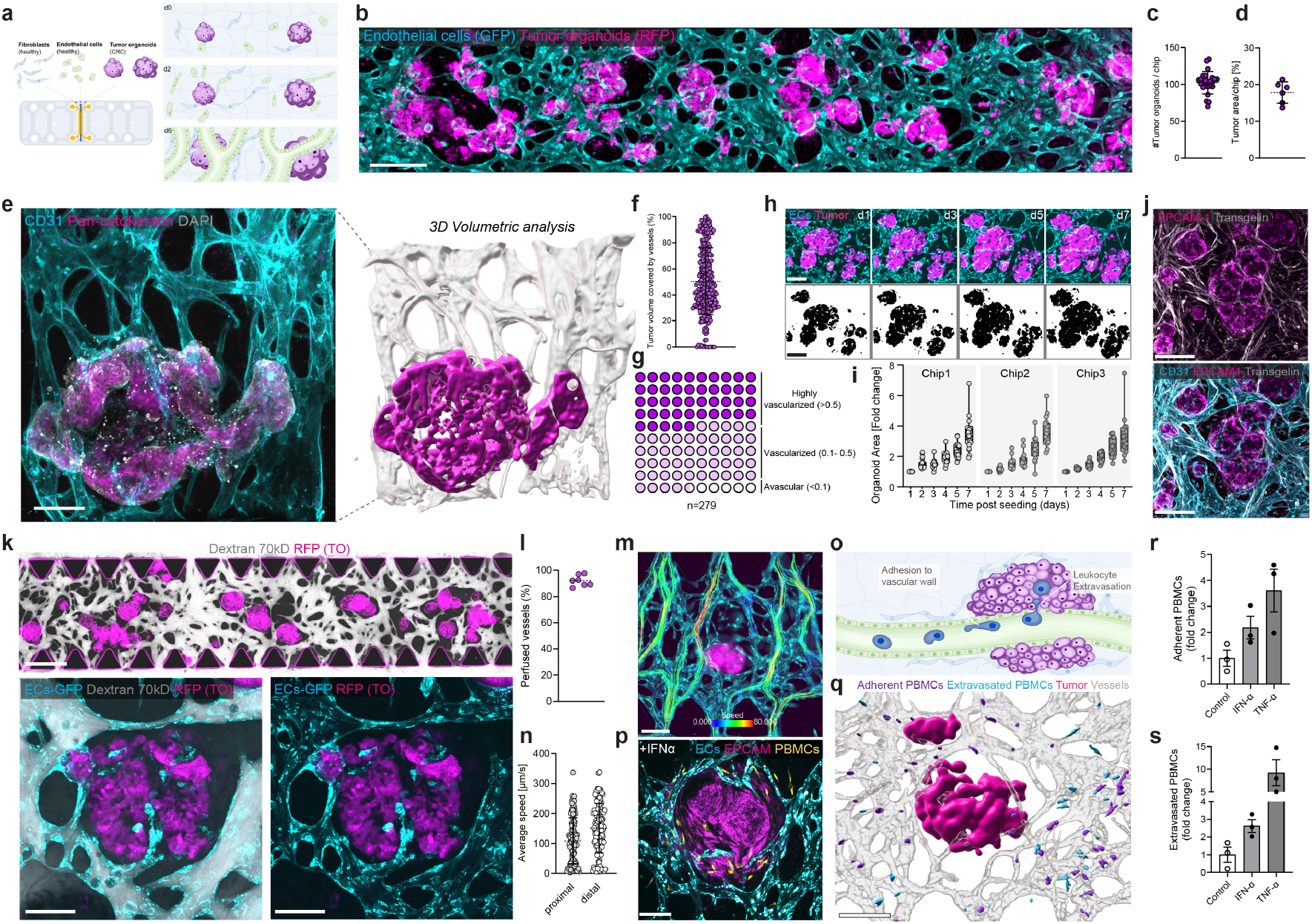
A perfusable vascularized tumor-stroma niche supports organoid growth and immune-vascular function. a, Schematic of VascO chip. Primary fibroblasts, endothelial cells and patient-derived tumor organoids are co-seeded and self-assemble into a vascularized tumor niche. b, Tile-scan confocal image of vascularized tumor organoids on day 9 after seeding (Endothelial cells GFP, cyan, Tumor organoids RFP, magenta).Scale bar, 500µm. c,d, Quantification of tumor burden per chip: number of organoids per chip (c) and tumor area fraction (d). Each point is one chip; bars denote mean ± s.d. e, Left, high-magnification confocal of a single tumor organoids (Pan-cytokeratin, magenta) surrounded by vessels (CD31, cyan) with tumor epithelium (pan-cytokeratin, magenta) and nuclei (DAPI). Right, 3D surface rendering of the same niche (vessels, grey; tumor, magenta).Scale bar, 100µm. f, Distribution of tumor–vessel contact/coverage scores across individual organoids. g, Classification of organoids by vascularization score (highly vascularized, vascularized, avascular); n = 279 organoids analysed. h, Longitudinal imaging of identical fields (days 1–7): representative confocal images (top) and binary segmentations (bottom). Scale bar, 200µm.i, Organoid growth kinetics expressed as fold-change in area relative to day 1 for three independent chips. Each point is one organoid; violins show distribution across organoids for 3 individual chips. j, Immunofluorescence demonstrating stromal organization: EPCAM1 tumor epithelium (magenta) in close apposition to TAGLN fibroblasts (transgelin; white) and CD31 vessels (cyan). Scale bar, 200µm. k, Functional perfusion of the vascular bed with 70-kDa fluorescent dextran (grey) confirms lumen continuity around tumor organoids; higher magnification inserts (bottom). Scale bar upper panel, 500µm, lower panels, 100µm. l, Fraction of perfused vessels per chip. n=7 entire chips from 3 biological replicates. m, Flow-velocity map of PBMCs (false-colour scale) perfused through vessels (GFP, cyan) surrounding a tumor organoid (RFP, magenta). Scale bar 200 µm. n, Quantification of mean flow speed in proximal versus distal vessel segments; each point denotes one PBMC. o, Schematic of leukocyte adhesion and extravasation assay on-chip. p, Immunofluorescence image of a representative niche after 24 h IFN-α stimulation. PBMCs (Cell tracker, yellow) perfused through the vessels (GFP, cyan) have engaged the tumor organoid (RFP, magenta). Scale bar, 100 µm. q, 3D rendering of the assay readout: adherent PBMCs (violet) on vessel walls and extravasated PBMCs (light blue) within the interstitium relative to vessels (grey) and tumor organoids (magenta). Scale bar, 200 µm. r,s, Quantification of adherent (r) and extravasated (s) PBMCs (fold-change over control) under control conditions and after pro-inflammatory stimulation (IFN-α or TNF-α). Bars, mean ± s.d.; dots represent independent chips.

Self-directed vascularization endows organoids with a perfusable microvasculature enabling immune cell trafficking, stromal interaction and clinically-relevant intravascular drug delivery. In practice however, the lack of spatiotemporal control inherent to self-assembly can lead to inefficient vascularization, reduced vessels-tumor interaction and poor functional integration. To overcome those challenges and establish a tractable platform amenable to the assessment of therapeutics, we established a method that promotes extensive vascular-tumor epithelial interface and enable direct vessel-tumor crosstalk by combining: (i) complete Matrigel depletion followed by single-cell clearance to generate scaffold-free, intact organoids that support vascular integration, (ii) an ultra-soft, engineered fibrin matrix that counters vessel thinning in the presence of tumor organoids, preserves lumen caliber, and sustains perfusability in co-culture, and (iii) a temporally staged VEGF-A + FGF-2 regimen during the initial vascularization window to enhance network maturation and connectivity (Fig. 1a). Confocal microscopy confirmed the formation of extensive endothelial networks closely interacting with RFP-expressing patient-derived tumor organoids and demonstrated robust and reproducible homogeneous tumor density between chips (Fig. 1b) and a high tumor load ( 100 organoids/chip, 18% of tumor area/chip; Fig. 1c,d). Three-dimensional volumetric analysis of pan-cytokeratin-stained tumor organoids confirmed extensive tumor-vascular interactions (Fig. 1e,f), with a significant proportion of tumor organoids exhibiting high vascularization levels (>0.5 volume ratio; Fig. 1g). Segmentation-based quantification over 7 days revealed reproducible 2 to 6 fold expansion of tumor organoid area despite withdrawal of Matrigel and canonical niche factors (Wnt3a/RSPO1/Noggin), indicating that the stromal compartment alone establishes a functional tumor-supportive niche (Fig. 1h,i). Immunofluorescence revealed supporting stromal fibroblasts (transgelin+) tightly interacting with tumor organoids (EPCAM1+) and aligned along endothelial structures (CD31+), frequently ensheathing vessels during self assembly (Fig. 1j and Supplementary Video 1,2). Perfusion with fluorescently-labelled Dextran (70 kDa) confirmed vascular network functionality (Fig. 1k), with a high degree of perfusable vessels (>95%) (Fig. 1l), demonstrating robust perfusion even in tumor organoid adjacent vessels (Fig. 1k). The circulation of peripheral blood mononuclear cells (PBMCs) confirmed that endothelial cells self organized into a functional perfusable vascular network with a uniform flow profile in proximal and distal tumor vessels (Fig. 1m,n and Supplementary Video 3,4,5). Treatment with IFN-α, used in clinic as a potent cancer immunotherapy (30), or the pro-inflammatory cytokine TNFa, for 24 h in the presence of perfused PBMCs, led to significant upregulation of adhesion molecules, immune cell adhesion and extravasation, highlighting the model’s capacity to accurately reproduce the vascular priming and immune trafficking central to IFN-α and TNFa activity (Fig. 1o-s, Extended Data Fig. 1 and Supplementary Video 6). This model provides undirected, self-assembled organoid vascularization, achieving functional tumor–stroma integration that supports tumoroid growth, sustains perfusion, enables immunecell trafficking, and responds to immune activation; hereafter referred to as VascO.

Solid tumors remodel their microenvironment to sustain growth, converting quiescent fibroblasts into cancer-associated fibroblasts (CAFs) and reprogramming endothelium into abnormal, leaky vasculature (31, 32). To define how patient tumors shape surrounding cell phenotype and function in VascO, we performed single-cell RNA sequencing on the VascO niche built with tumor organoids from three donors and matched healthy vasculature as control (Fig. 2a). To enable efficient tissue dissociation for downstream single-cell profiling, we developed a novel two-step protocol that first selectively depletes stromal cells from the anastomosed media channels flanking the 3D tissue, followed by incubation with a Nattokinase-based enzymatic cocktail (Fig. 2a). Libraries were generated 9 days post-seeding, when vascular networks had matured into perfusable vessels (established by day 5) and cultures were maintained in growth factor-reduced medium to promote vessel stabilization and sustain physiological cell–cell interactions. Heterogeneity analysis and visualization using uniform manifold approximation and projection (UMAP) embedding of all cells resolved ten major clusters: four fibroblasts, two endothelial, and four epithelial states across all three donors (Fig. 2b-d). Unsupervised hierarchical clustering within fibroblasts revealed four transcriptional programs, highlighting populations harboring markers associated with oxidative stress response (Cluster 1), extracellular matrix (ECM) production and myofibroblasts (Cluster 2), proliferation (Cluster3), and inflammation (Cluster 4) (Fig. 2e). The top differentially expressed genes for each cluster aligned with canonical CAFs archetypes. Cluster 2 expression of ECM-related genes such as COL1A1/1A2/3A1/8A1/15A1/16A1, FN1 and the myofibroblasts-related genes ACTA2, ACTG2, FAP, POSTN, TAGLN, TGFB2, WNT5A indicative of contractile and strongly activated cells with a role in ECM production and matrix stiffening, closely overlapped with key known markers of myofibroblastic CAFs (myCAFs) (33–38). Cluster 3 signature highlighted markers indicative of a high mitotic activity including MKI67, TUBA1B, CCNB1/2, CDK1, PLK1, reminiscent of a dividing CAFs (dCAFs) phenotype (39), while cluster 4 highest differentially expressed genes IL6, CXCL1/2/3/6/8/12, C3, SOD2, PDGFRa, STAT3, CCL2, and ICAM1 revealed pro-inflammatory markers consistent with the transcriptional profile of inflammatory CAFs (iCAFs) (33, 34, 40–43). Gene-ontology analysis of fibroblasts co-cultured with donor-derived tumoroids revealed upregulation of angiogenic (sprouting angiogenesis, endothelial migration, NOTCH), adhesion/motility and collagen-rich ECM programs, with concomitant suppression of oxidative-phosphorylation/mitochondrial activity, consistent with a shift from quiescence to a glycolytic activated CAF-like state (Fig. 2f). Comparing fibroblast transcriptomes between tumor co-culture and healthy-vasculature controls identified six programs differentially modulated by tumor organoids (Fig 2g). Across all donors, tumoroids strongly suppressed fibroblasts proliferation, as evidenced by the downregulation of markers such as MKI67, CDC20 or CCNB1, and consistently induced myofibroblasts (ACTA2, ACTG2, TAGLN, FAP, MYH9 and 10) and angiogenesis (VEGFA, ANGPTL2, PDGFA, PDGFRA) signatures, thus emphasizing the plasticity of healthy human primary fibroblast and their ability to acquire CAFs signatures when cultured in close proximity to tumor organoids (Fig. 2g). Importantly, inflammatory markers (IL6, CXCL1/2/3/8) revealed donor-specific immune states, with donors 2 and 3 driving a pro-inflammatory phenotype and donor 1 promoting an anti-inflammatory microenvironment, concordant with lower cytokine levels in circulating media (Fig. 2h). ECM-related signatures (COL1A1/1A2/3A1/5A1/6A3) were also donor-dependent (Fig. 2g) and confirmed by immunofluorescence analysis (Fig. 2i,j). Imaging showed FAP fibroblasts forming a peritumoral cuff and perivascular stromal strands aligned with vessels, recapitulating the spatial heterogeneity of FAP CAFs (44, 45) (Extended Data Fig. 2a). Together, these findings demonstrate that patient-derived tumor organoids instruct human fibroblasts to adopt a CAF-like state, generating a donor-specific, tumor-educated microenvironment within the VascO niche.

**Figure 2.**
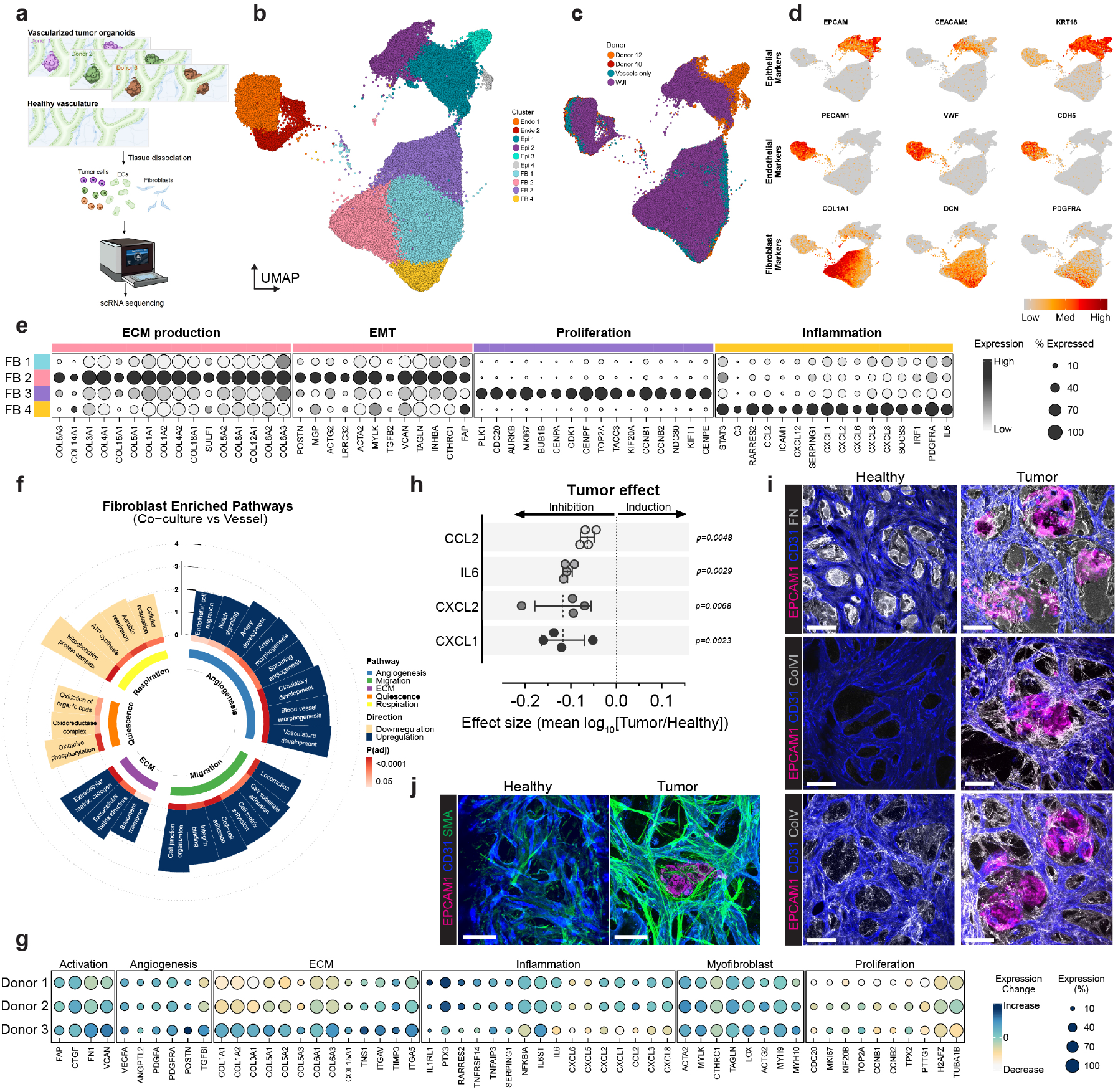
Single-cell analysis reveals tumor-driven fibroblast states and donor-dependent inflammatory programs in a vascularized organoid niche. a, Experimental design. Vascularized tumor organoids were generated from three patient donors and compared with matched healthy vasculature controls. Chips were enzymatically dissociated and profiled by single-cell RNA-seq. b, UMAP of all cells coloured by major clusters resolves epithelial (Epi 1–4), endothelial (Endo 1–2) and fibroblast (FB 1–4) populations. c, UMAP colored by sample of origin (donors 1, 2, 3 and vasculature-only control), showing broad representation of each state across donors. d, Feature plots of canonical markers used to assign identities: epithelial (EPCAM, CEACAM5, KRT18), endothelial (PECAM1, VWF, CDH5) and fibroblast (COL1A1, DCN, PDGFRA). Signal intensity denotes scaled expression. e, Dot-plot of fibroblast subclusters (FB1–FB4) highlighting programs related to extracellular-matrix (ECM) production, epithelial–mesenchymal transition (EMT), proliferation and inflammation. Dot size indicates the percentage of cells expressing the gene; color indicates average expression per cluster. f, Gene-set enrichment analysis of fibroblasts in tumor co-culture versus vessel-only cultures. Pathways are grouped into major functional categories (angiogenesis, migration, ECM, quiescence and respiration). Inner ring colour indicates pathway category; blue and yellow bar color indicate the direction of regulation (upregulation or downregulation); and colour intensity of the red inner bar corresponds to the adjusted P value. g, Tumor-induced modulation of fibroblast programs across the three donors. Relative to healthy vasculature, tumor organoids co-culture suppresses proliferation while inducing activation/contractile, pro-angiogenic and ECM genes, with donor-dependent inflammatory and ECM signatures. Color encodes direction and magnitude of expression change; dot size reflects the percentage expressing. h, Tumor effect sizes for secreted inflammatory mediators in tumor organoid versus healthy conditions (mean log10[Tumour/Healthy]); points denote individual chip, bars indicate mean effect; p-values from two-sided tests as indicated. i, Confocal imaging of healthy versus tumor conditions showing EPCAM1 epithelium (magenta), αSMA stromal fibres (green) and CD31 vessels (blue); tumor niches display dense, peritumoral stroma closely apposed to vessels.Scale bar, 200 µm. j, Immunofluorescence of EPCAM1 (magenta), CD31 (blue) and ECM proteins (fibronectin, FN; collagen IV, ColV; grey) demonstrating remodelling of the peritumoral matrix in tumor niches compared with healthy controls. Scale bar, 100 µm.

The impact of tumor organoids on healthy endothelial cell fate was investigated through transcriptional and functional analyses of tumor-associated vessels. Single cell RNA sequencing identified two distinct clusters of endothelial cell transcriptional responses (Endo1 and Endo2). Endo1 exhibited an angiogenic/tip-like program enriched for migration and guidance receptors (CXCR4, UNC5B, ACKR3), whereas Endo2 comprised cycling endothelium expressing core G2/M–S-phase regulators (MKI67, CDC20, CCNA2, CDK1, CCNB2) (Fig. 3a). Further analysis highlighted a downregulation of endothelial barrier (KLF2, JAM3) and inflammatory genes (e.g. CXCL1/2/5/8) in the presence of tumor organoids as well as donor-dependent differential expression patterns in cell adhesion and extracellular matrix remodeling (Fig. 3b). Angiogenesis-associated genes, particularly those linked to tip-cell fate such as ESM1, ANGPT2, CXCR4 and EFNB2 exhibited marked induction upon co-culture with tumor organoids demonstrating tumor-driven endothelial reprogramming (Fig. 3c). Microscopic analysis of endothelial morphology confirmed that tumor vessels displayed an irregular and disorganized pattern and showed excessive CD31+ filopodia formation and vessel sprouting, in contrast to cultures without tumor tissue (Fig. 3d and Extended Data Fig. 2b). Quantitative morphometric analyses further revealed abnormal and excessive vascular branching in the peritumoral region, reflected by elevated counts of vessels branch points (385.8 vs 82.3; p=0.0065), junctions (218.7 vs 50.9; p=0.0074), and terminal ends (106.7 vs 11.5; p=0.0061) (Fig. 3e). It is widely recognized that tumor-driven angiogenesis generates immature, disorganized vascular networks, resulting in hyperpermeable vessels (46). To assess the functional consequences of aberrant tumor vasculature in our model, we perfused fluorescently labelled dextran (70kD) through the endothelial networks, and indeed observed a marked increase in vascular permeability in the presence of tumor organoids (Fig. 3f and Supplementary Video 7,8). As a positive control, treatment with recombinant VEGF induced a dose-dependent increase in permeability, confirming vessels sensitivity to canonical permeability drivers (Extended Data Fig. 3). As VEGF-A and Angiopoietin-2 (Ang2) are key drivers of vascular leakage (47, 48) we treated the co-cultures with a bispecific anti–VEGF-A/anti–Ang2 antibody and assessed vessel permeability. Blocking VEGF/Ang2 for 24 h significantly reduced tumor-induced vascular leakage (Fig. 3g).

**Figure 3.**
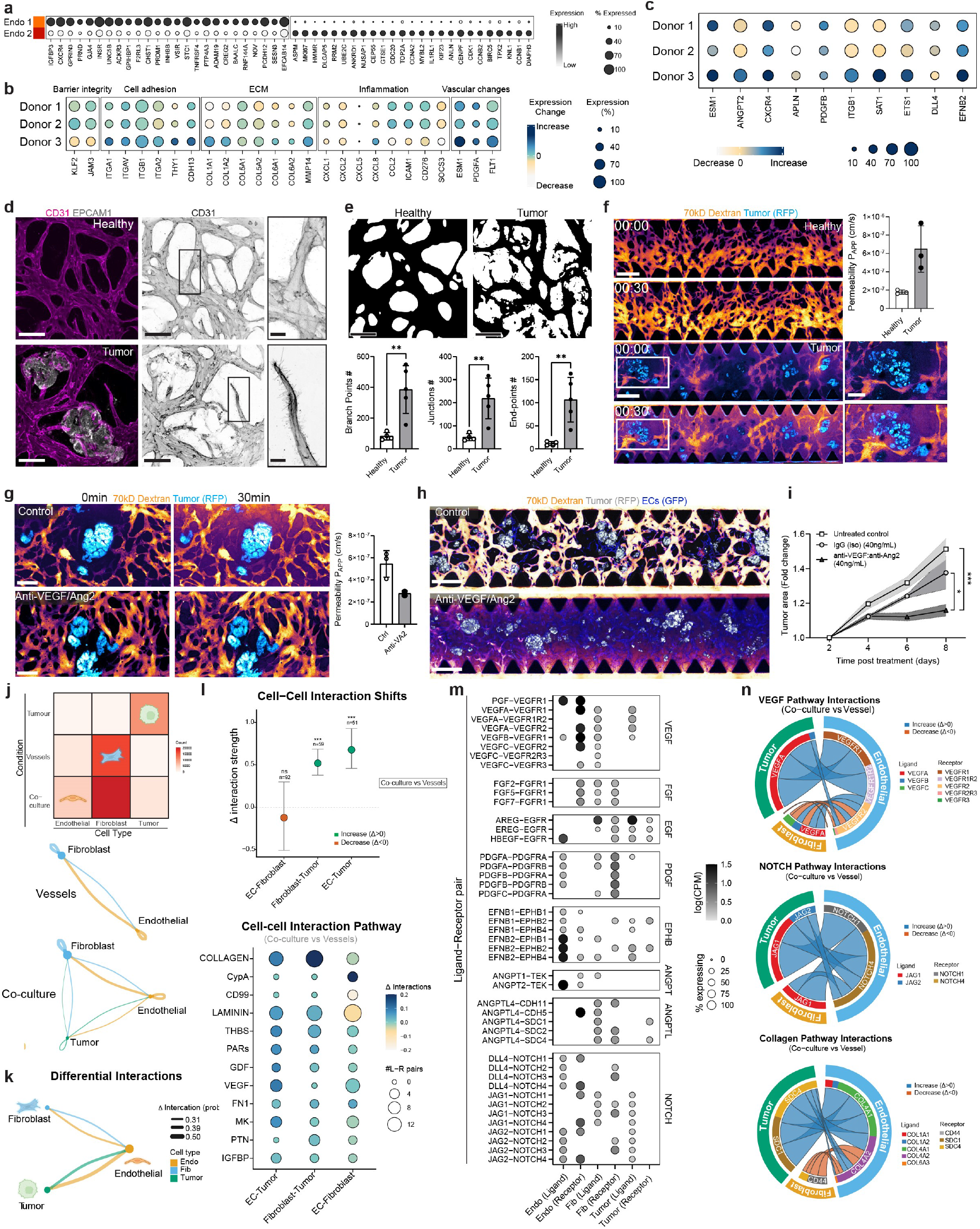
Vascularized tumor organoids reprogram endothelial cells by increasing tip cell fate and VEGF/Ang2-driven leakiness. a, Dot plot of top differentially expressed genes from single-cell RNA sequencing of endothelial cells co-cultured with tumor organoids, identifying an angiogenic/tip-like cluster (Endo1) and a proliferating cluster (Endo2); b, Dot plot illustrating changes in gene expression grouped by biological function after co-culture with tumor organoids. Dot size shows the proportion of expressing cells; colour encodes direction and magnitude of change (co-culture versus vessels alone). c, Dot plot showing marked upregulation of tip-cell associated genes in endothelial cells upon co-culture with tumor organoids from three different donors. d, Representative 3D immunofluorescence images of endothelial networks in healthy versus tumor co-culture conditions. Vessels are stained for CD31 (magenta) and tumor organoids are visualized by EPCAM1 (white). Tumor co-cultures exhibit disorganized vessel morphology with excessive filopodia (insets).Scale bar, 100 µm, insert 20 µm. e, Morphometric analysis of vascular networks. Data are presented as mean ± s.d. Scale bar, 100 µm. f, Assessment of vascular permeability. Time-lapse images show leakage of perfused 70kD Dextran (orange) in healthy versus tumor (blue) co-cultures over 30 minutes. Quantification of the apparent permeability coefficient (Papp) demonstrates significantly increased leakage in tumor-associated vessels. Data are presented as mean ± s.d. Scale bar, 500 µm, insert 200 µm. g, Dual VEGF/Ang2 blockade reduces dextran extravasation. Co-cultures were treated with a bispecific anti–VEGF-A/Ang2 antibody for 24h. Representative images and quantification show a significant reduction in tumor-induced vascular leakage upon treatment. Data are presented as mean ± s.d. Scale bar, 200 µm. h, Representative images showing the effect of sustained anti–VEGF-A/Ang2 treatment on vascular network formation in the presence of GFP-expressing tumor organoids (white). i, Longitudinal quantification of tumor organoid growth over 8 days. Fold change in tumor area is plotted for untreated controls (UTC), IgG isotype control, and anti–VEGF-A/Ang2 treated co-cultures. Data are presented as mean ± s.e.m. Scale bar, 500 µm. j, Cell–cell communication inferred from single-cell transcriptomes. Top, interaction matrix summarising the frequency/strength of significant ligand–receptor (L–R) pairs between cell types. Bottom, network views for vessels alone and tumor co-culture; edge width reflects interaction strength/number of significant L–R pairs. k, Differential interaction network highlighting gains (orange) and losses (blue) in L–R communication in tumor co-culture relative to vessels alone. l, Aggregate shifts in interaction strength by communicating pairs (for example EC–tumor, EC–fibroblast, fibroblast–tumor). Bottom, bubble heat map of pathway-level changes; circle area reflects the number of significant L–R pairs and colour reflects the direction of change. m, Selected ligand–receptor families showing differential usage across cell pairs: VEGF/VEGFR, FGF/FGFR, EGFR ligands, PDGF/PDGFR, ephrin/EPH, ANGPT/TIE, integrins and NOTCH. Dot size represents frequency of significant L–R pairs; shading indicates relative expression/enrichment. n, Circos-style pathway summaries for VEGF, NOTCH and collagen signalling showing sender (ligand-producing) and receiver (receptor-expressing) cell types. Segment colour denotes pathway; ribbon thickness indicates interaction load; inner ring indicates increase (up) or decrease (down) in co-culture versus vessels alone.

Sustained treatment with the anti-VEGF/anti-Ang2 bispecific antibody, originally developed to suppress pathological angiogenesis and cancer progression (49), also prevented the establishment of functional, perfusable vascular vessels in our model (Fig. 3h). Importantly, longitudinal imaging of RFP-marked tumor organoids demonstrated substantial suppression of tumor growth over 8 days in culture in the presence of Anti VEGF/anti-Ang2, demonstrating that vascularization actively supported tumor organoid expansion (Fig. 3i). Cell-cell communication (ligand-receptor) analysis inferred from single-cell transcriptomes indicated that tumor organoids shift stromal communication networks towards a tumor-centered niche, strengthening tumor-endothelial signaling while reducing endothelial-fibroblast interactions (Fig. 3j,k). Pathway-level analysis of inferred ligand-receptor interactions showed coordinated increases in VEGF, NOTCH and collagen-rich ECM exchange at the vessel interface (Fig. 3l), underpinned by canonical ligand-receptor pairs such as VEGFA-VEGFRs, JAG1/2-NOTCH1/4 and COL1A1-CD44/SDC receptors (Fig. 3m,n). Together, these findings indicate that patient-derived tumor organoids can actively reshape endothelial cell phenotypes, driving angiogenic remodeling and increasing vascular permeability predominantly via VEGF/Ang2 signaling.

Cancer immunotherapies targeting solid tumors frequently face dose-limiting toxicities, as many tumor-associated antigens are also expressed in healthy tissues, thereby attenuating their clinical efficacy. To address these limitations, next-generation tumor-restricted therapeutics have been engineered to minimize on-target, off-tumor effects and improve therapeutic index (50). Yet, by engaging increasingly complex and human-specific immune mechanisms (50), these inrnovative approaches expose the limitations of conventional preclinical models. To assess whether VascO could be used to support the assessment of new tumor-activated immunooncology drug candidates, we started by testing the platform ability to assess the efficacy and safety profile of a first generation CEACAM5-targeting T cell-bispecific (TCB) antibody. CEACAM5-TCB is an IgG1-based bispecific antibody comprising a heterodimeric anti-CD3ε arm and two anti-CEACAM5 arms, designed to redirect T cells toward CEA-expressing tumor cells (51, 52) (Fig. 4a). CEACAM5-TCB treatment at clinically relevant concentrations (53) for 72 h induced robust recruitment, extravasation, and engagement of perfused immune cells with tumor organoids as demonstrated by high-resolution 3D imaging and segmentation of whole tissues (Fig. 4b-e and Supplementary Video 9), leading to localized apoptosis as evidenced by increased cleaved Caspase-3 in vascularized tumor organoids compared with non-treated control (Fig. 4f). To assess the safety profile of CEACAM5-TCB, we generated donor-matched, vascularized healthy-colon organoids from tumor-adjacent tissue and reduced thrombin concentration to yield larger-diameter vessels with sustained perfusion (Extended Data Fig. 4, Supplementary Video 10). Treatment of healthy vascularized donors matched colon organoids with clinical relevant doses of CEACM5-TCB showed comparable immune infiltration and organoid killing (Fig. 4g-j). These findings are consistent with the severe intestinal adverse events observed in clinical trials (54, 55) and the lack of therapeutic window leading to the internal decision to terminate the program. To mitigate on-target, off-tumor toxicity, a novel protease-activated CEACAM5-TCB (Prot-CEACAM5-TCB) was engineered, in which the anti-CD3 arm is masked by an anti-idiotypic scFv tethered via a protease-cleavable peptide linker (Fig. 4k). This strategy enables selective activation within the tumor microenvironment, where protease activity removes the mask and restores TCB potency (56). In the tumor VascO model, Prot-CEACAM5-TCB, upon proteolytic unmasking, induced immune-cell infiltration and tumor cell killing comparable to the unmasked CEACAM5-TCB, indicating that conditional activation preserves intrinsic potency (Fig. 4l,m). However, in sharp contrast to CEACAM5-TCB, Prot-CEACAM5-TCB remained minimally active up to 10 µg/mL in matched healthy VascO cultures, demonstrating tumor-restricted engagement (Fig. 4n,o). Across paired tumor/healthy VascO, full dose-response analyses showed that Prot-CEACAM5-TCB expands the in vitro therapeutic window, increasing tumor-specific cytotoxicity while suppressing off-tissue effects, thereby illustrating how the VascO platform can resolve selectivity and quantify therapeutic index for conditionally activated bispecifics (Fig. 4p-r). These results establish VascO as a high-resolution platform to evaluate efficacy and safety profiles of next-generation immunotherapies.

**Figure 4.**
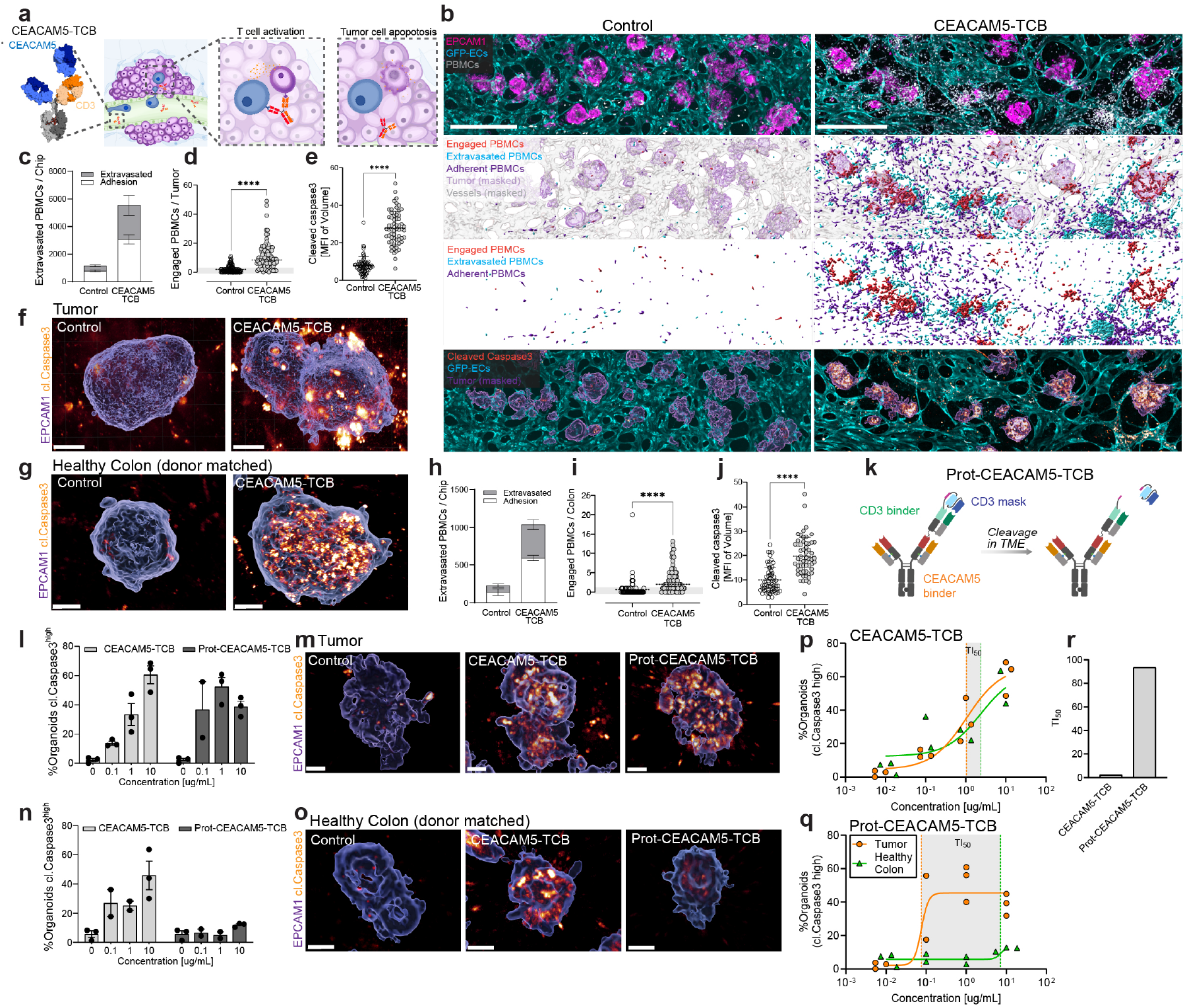
Vascularized, donor-matched tumor and healthy organoids enable therapeutic window assessment of novel immuno-oncology drugs. a, Schematic illustrating the mechanism of action for the CEACAM5 T-cell bispecific antibody (TCB). The antibody cross-links the CD3 receptor on T-cells with the CEACAM5 antigen on tumor cells, triggering T-cell activation and subsequent tumor cell lysis. b, Representative 3D confocal projections of tumor co-cultures under vehicle control (left) or CEACAM5–TCB (right). EPCAM+ epithelium (magenta), GFP+ endothelium (cyan) and PBMCs (white) are shown. Middle, computational masks classify PBMCs as adherent (vessel-associated), extravasated (outside vessels) or engaged (in contact with tumor). Bottom, cleaved caspase-3 (orange) marks apoptotic tumor cells. Scale bar, 500 µm. c–e, Quantification of vascular trafficking and killing in vascularized tumors: c, quantification of extravasated and adherent PBMCs; d, number of engaged PBMCs per tumor organoid; e, apoptotic burden (cleaved caspase-3 events/area). CEACAM5–TCB markedly increases PBMC extravasation and engagement and induces tumor apoptosis. Data are presented as mean ± s.d. f, Representative confocal images of vascularized tumor organoids (EPCAM+, blue) show widespread caspase-3 activation (orange) after CEACAM5–TCB compared with control. Scale bar, 50 µm. g, Donor-matched vascularized healthy colon organoids (EPCAM+, blue) exhibit strong on-target killing (orange) with CEACAM5–TCB. Scale bar, 50 µm. h–j, Quantification in healthy colon: h, number of extravasated and adherent PBMC; i, engaged PBMCs per colon organoid; j, apoptotic burden. CEACAM5–TCB enhances PBMC trafficking and killing in healthy colon. k, Design of a protease-activatable CEACAM5–TCB (Prot-CEACAM5–TCB) in which the CD3-binding arm is masked and released by proteases abundant in the tumor microenvironment (TME), restricting activity to tumors. l, m, Tumor efficacy of CEACAM5–TCB versus Prot-CEACAM5–TCB. l, Percentage of tumor organoids positive for cleaved caspase-3 after 72h across antibody dose-response; m, representative images. The protease-activatable format retains tumor-killing potency. Scale bar, 50 µm. n, o, Healthy colon toxicity with CEACAM5–TCB versus Prot-CEACAM5–TCB. n, Percentage of colon organoids with caspase-3 activation across dose-response; o, representative images. Protease activation strongly reduces killing of healthy colon. Scale bar, 50 µm. p, q, Dose–response curves for tumor (orange circle symbols) and donor-matched healthy colon (green triangle symbols) treated with p, CEACAM5–TCB or q, Prot-CEACAM5–TCB. Curves show non-linear fits; vertical dashed lines mark the concentrations used to compute the therapeutic index (TI50). r, Summary of TI50 (ratio of the concentration producing 50% caspase-3 positivity in healthy colon to that producing 50% killing in tumor). Prot-CEACAM5–TCB exhibits a substantially higher therapeutic window than CEACAM5–TCB.

We thus describe a vascularized tumor organoid system comprising patient-derived tumoroids vascularized with a functional capillary network supported by fibroblasts, that we used for the assessment of the efficacy and safety of novel cancer immunotherapy. We show that tumor cells reprogram neighboring fibroblasts and endothelial cells toward cancer-associated phenotypes. The magnitude of this reprogramming is patient-specific and manifests as phenotypic and functional changes, including altered vascular permeability and capillary-network architecture. Reciprocally, vascularization supports tumoroid growth, recapitulating the well-described crosstalk between tumors and the surrounding vasculature (32). By leveraging the functional vascular compartment and perfusing circulating immune cells, we could validate the design of a next-generation tumor-activated TCB by demonstrating its maintained efficacy in killing tumor cells and risk-benefit superiority in sparing healthy intestinal tissue, thus demonstrating the applicability of VascO for preclinical efficacy and safety assessment in drug development.

VascO uniquely captures and enables perturbation of the transitional window in which naïve human stroma self-organizes into a tumor-conditioned vascular-fibroblast niche, enabling causal dissection of events that patient biopsies rarely resolve. This window encompasses the angiogenic switch, a decisive early step in tumorigenesis, and the ensuing conversion of endothelium into immature, hyperpermeable networks driven by VEGF-A and reinforced by Ang2, processes long implicated in leak, hypoxia, and immune exclusion (57). Because VascO is perfusable, these endothelial transitions can be followed longitudinally and physiologically challenged with normalization strategies like dual VEGF-A/Ang2 blockade that prolongs normalization and boosts antitumor immunity in vivo, linking mechanism to measurable changes in leak, perfusion, and immune trafficking. In parallel, VascO reveals fibroblast bifurcation into canonical myCAF/iCAF-like states and allows testing of microenvironmental drivers shown by multi-omics to organize CAF neigh-borhoods across cancers (58, 59). The convergence of time-resolved angiogenic remodeling with CAF plasticity in a human setting creates a tractable platform to uncover early or-ganizing principles of the tumor microenvironment and to ask whether vascular normalization can reprogram stromal states to favor immunity, an idea supported by preclinical work but not mechanistically mapped in human tissues (60). Finally, because patient-derived organoids predict clinical drug sensitivity (19–22), embedding them in a self-assembled stromal-vascular unit turns VascO into a discovery engine for new biology with direct translational read-through.

In the meantime, regulatory agencies worldwide are rapidly refining their frameworks to promote alternative methods, with mounting initiatives underway to reduce, and even selectively phase out the use of animal models in drug development and broader applications within the next decade (14, 15, 61–65). Achieving this ambitious goal is contingent upon the establishment of highly predictive human-relevant systems that can improve on current preclinical tools. Among therapeutic areas, cancer, and particularly immuno-oncology, is poised to lead this transition given its life-threatening nature, the limited translational value of animal models, the urgent unmet medical need, and the growing oncology regulatory filings that rely on in vitro approaches (66). VascO directly addresses this challenge by enabling the vascularization of virtually any tissue-derived organoid, thereby extending applications well beyond colorectal cancer. As such, VascO represents an enabling technology for the future tool box of animal-free drug development by providing a human-relevant, mechanistically rich platform that is consistent with emerging regulatory expectations and can help meet the rising demand for predictive and ethical alternatives in cancer immunotherapy and beyond.

## ACKNOWLEDGEMENTS

We thank Anna Krohmer-Rombach for creating the illustrations, Antonello De Bari and Jennifer Hinke for technical support.

## AUTHOR CONTRIBUTIONS

R.W. and R.V. conceived the study; R.W. and R.V. wrote the manuscript; R.W. and R.K. developed the protocol and optimized culture conditions; R.W., R.K. and L.H. designed and performed most experiments in the manuscript; N.G. isolated and supplied organoid lines from primary tissue; I.C. and F.K. performed scRNAseq experiments and generated sequencing libraries; F.M. and S.N. performed the computational analysis of scRNAseq and data interpretation was performed by R.W. and R.V.; R.W. performed immunohistochemistry stainings, imaging and analysis with support from R.K., L.H., N.L. and T.K.; Live imaging videos were generated by R.W. with the support of R.K.; I.W., C.F.K. and M.B. provided the T-cell bispecific molecules and supported with data interpretation of the pharmacology studies; All authors read and approved the manuscript

## DATA AVAILABILITY

Raw and processed single cell data with cell type annotation are available upon request. Additional data and materials are available from the corresponding author upon reasonable request, subject to any restrictions imposed by ethics approvals and material transfer agreements.

## Methods

### Cell Culture

All cells were maintained in a humidified incubator set to 37°C and 5% CO_2_. Culture media was replaced every two days. Upon reaching confluence, cells were washed twice with 1 × DPBS (Gibco) devoid of calcium and magnesium, then detached using pre-warmed Trypsin-EDTA (0.05%) phenol red (Gibco). Human umbilical vein endothelial cells (HUVECs) (Angio-Proteomie) expressing GFP were cultured in EGM-2 (Lonza) up to passage 8. Human liver sinusoidal microvascular endothelial cells (LSMVECs) (Angio-Proteomie) expressing GFP were cultured in EGM-2 MV (Lonza) up to passage 9. Normal human lung fibroblasts (NHLF) (Lonza) were cultured in FGM-2 (Lonza) up to passage 7.

### Human samples and Ethics statement

Human intestinal tissue resections, along with concurrent data collection and experimental procedures, were conducted within the framework of the non-profit foundation HTCR (Munich, Germany), including informed patient consent. The HTCR Foundation’s framework received approval from the ethics commission of the Faculty of Medicine in the Ludwig Maximilian University (no. 025-12) and the Bavarian State Medical Association (no. 11142). We used micro- and/or macroscopically tumour-free regions of resectates for further preparation and downstream handling.

### Organoids culture

Human colorectal cancer organoids and adjacent healthy colon organoids were obtained as previously described (). In short, after isolating intestinal crypts using the previously described protocol (StemCell Technologies), the crypts were embedded in a 25 µl dome of growth-factor-reduced Matrigel (Corning). The droplet containing organoids was then cultured in a 50% (v/v) mix of Human IntestiCult OGM human basal medium (OGM; StemCell Technologies) and organoid supplement (StemCell Technologies) in a 24-well clear TC-treated plate (Corning Costar). After seeding, the media was supplemented with Y27632 (10 µM l^−1^; StemCell Technologies) and 1% penicillin/streptomycin (Gibco). Throughout the week of expansion, OGM was changed every 2 days, deprived of Y27632. For further storage or experimental use, organoids were either kept in culture or frozen in CryoStor CS10 (StemCell Technologies) and stored in liquid nitrogen.

For organoid maintenance, cryopreserved organoid vials were thawed, and the content of a vial was then transferred into of 1% BSA solution at 4°C. The solution was centrifuged for 4 min at 250g at 4°C. The supernatant was discarded, and the pellet was resuspended in 25 µl Matrigel domes on a 24-well plate. Finally, IntestiCult OGM Human Basal Medium (Corning) supplemented with Y27632 (StemCell Technologies) was added, and plates were incubated at 37°C, 5% CO_2_. For splitting, medium was aspirated, and wells were washed once with PBS at room temperature. Then, Gentle Cell Dissociation Reagent (StemCell Technologies) was added, incubated at room temperature, and transferred to a 15 ml tube. The tube was centrifuged for 4 min at 250g at 4°C. Afterwards, the supernatant was discarded, and cell pellets were resuspended in cold 1% BSA solution. The cell solution was centrifuged for 4 min at 250g at 4°C. The cell pellet was resuspended in Matrigel and plated as previously described.

Before seeding organoids into the microfluidic chip, healthy organoids underwent a 3-day treatment with 1:1000 DAPT, followed by an additional 2 days in IntestiCult ODM Human Organoid Differentiation Medium (Corning). To harvest, organoids were washed once with cold PBS. Then, cold Cell Recovery Solution (Corning) was added, and the domes were gently scraped off the plate. The plate was incubated for 30–40 min on ice while shaking. Organoids were then transferred into a 15 ml Falcon tube and centrifuged at 300g for 5 min at 4°C. After centrifugation, organoids were washed with cold 1% BSA solution. The required amount of organoid solution for each experimental condition was aliquoted, centrifuged at 300g for 5 min, and stored on ice. Just prior to loading onto the chip, organoids were resuspended in the appropriate amount of cell mix in fibrinogen, mixed with thrombin, and then loaded onto the chip.

### ECM Preparation and Device Seeding for Vascularization of Organoids

Human fibrinogen (Sigma-Aldrich) was solubilized in sterile DPBS (Gibco) according to the manufacturer’s guidelines to a final concentration of 10 mg/ml. Following sterile filtration through a 0.22 µm filter, it was aliquoted and stored at − 80°C. Thrombin from human plasma (Sigma-Aldrich) was dissolved in sterile DPBS (Gibco) as per manufacturer’s instructions to a final concentration of 100 U/ml. After sterile filtration through a 0.22 µm filter, it was aliquoted and stored at − 20°C.

Prior to experimental use, fibrinogen and thrombin aliquots were thawed on ice. Thrombin was diluted to a working concentration of 50 U/ml. Endothelial cells and fibroblasts were detached as described above and centrifuged at 250g for 5 min. Following resuspension, cells were counted and combined in a 2:1 ratio (endothelial cells to fibroblasts). After a subsequent centrifugation (250g, 5 min), cells were resuspended in cold fibrinogen and mixed with either healthy or tumor organoids.

The cell suspension in fibrinogen was then mixed with thrombin to achieve a final thrombin concentration of 1 U/ml. Immediately after mixing, the cell suspension was loaded into the idenTx 3 chip (AIM Biotech) via the gel loading port. The microfluidic chips, placed in the idenTx holder (AIM Biotech), were transferred to a humidified incubator (37°C, 5% CO_2_) for 15 min to allow fibrin gel polymerization. Subsequently, 15 µl of fibronectin solution (60 µg/ml) was added to the side channels, and the chips were incubated in the humidified incubator (37°C, 5% CO_2_) for an additional 30 min. After this 30 min incubation, 70 µl of fresh, pre-warmed EGM-2 MV media was added to one side of the chip and 50 µl to the opposite side. Once the media equilibrated, it was aspirated and replaced with EGM-2 MV media supplemented with 50 ng/ml human vascular endothelial growth factor (VEGF) (Peprotech) and 50 ng/ml human fibroblast growth factor 2 (FGF-2) (R&D Systems) for the initial four days of culture. Media was replenished daily by aspirating old media and adding 70 µl of fresh media to one side and 50 µl to the opposite side. On the following day, the idenTx holders containing the microfluidic chips were transferred to the OrganoFlow rocker (Mimetas), oriented parallel to the rocker’s axis. The rocking interval was 4 h with an angle of 25°.

On the third day of culture, the side channels of the idenTx 3 chip were seeded with the same type of endothelial cells previously cultured in the gel. Endothelial cells were detached as described above and centrifuged at 250g for 5 min. Following resuspension in media, cells were counted and diluted to a concentration of 1.5 × 10^6^ cells/ml. Media was aspirated from all media ports on the idenTx3 chips. Then, 60 µl of media was added to two ports on the same side of the chip, and 50 µl of media was added to the opposite side. After the 50 µl addition, 10 µl of the cell suspension was added to the media ports that previously received 60 µl of media. The microfluidic chips were then incubated in a humidified incubator (37°C, 5% CO_2_) for 30 min. Following incubation, the media was exchanged as described previously, and the microfluidic chips were returned to the OrganoFlow rocker. A perfusable microvascular network was observed after 4 days.

### Treatment

### Immune cell preparation

Cryopreserved human peripheral blood mononuclear cells (PBMCs) were thawed and immediately diluted in pre-warmed media, followed by centrifugation and washing to remove freezing media. Unless specified otherwise, cells were centrifuged at 250g for 5 min. Afterwards, cells were maintained in T25 Ultra-Low Attachment Flasks (Corning) and incubated at 5% CO_2_ and 37°C with humidity for proliferation in PBMC culture medium (RPMI + Glutamax + 10% FBS + 1% MEM NEAA + 1% Sodium Pyruvate + 1% Pen/Strep). For cell trace staining, PBMCs were harvested, and the appropriate cell count was stained with CellTrace™ Far Red Cell Proliferation Kit (ThermoFisher Scientific) according to the supplier protocol and used for infusion onto the vascularized chip.

### Immune cell infusion

Following vascular network maturation, immune cell perfusion was performed via the side channels of the idenTx9 chip. The prior fluorescently labeled PBMCs resuspended, counted and diluted to desired concentration of 1.5 × 10^6^ cells/mL in EGM-2 MV : PBMC culture media (as described above). The cell suspension was introduced into the upper channel via a pressure gradient. Vascularized chips were then incubated at the OrganoFlow rocker (Mimetas), oriented parallel to the rocker’s axis. The rocking interval was 8 min with an angle of 25° in a humidified incubator (37°C, 5% CO_2_).

### Compound treatment

Vascularized microfluidic chips were treated with the indicated compounds prepared at defined concentrations in EGM-2 MV medium (Lonza). Treatments were introduced into the vascular channels using a pressure gradient to ensure uniform perfusion across the network. Culture medium was refreshed every 24 h throughout the treatment period.

### Dextran perfusion and permeability studies

Barrier function and permeability of the engineered microvascular networks was quantified using fluorescently labeled dextran as a tracer molecule, following established procedures (xx). Media containing 50 µg/mL 70 kDa-CF633 conjugated dextran (Biotinum #80141) was infused into the network and real-time confocal fluorescence imaging was performed using a Leica SP8 or Stellaris 8 confocal microscope. Images were analyzed using ImageJ. The apparent permeability was calculated using the following equation:

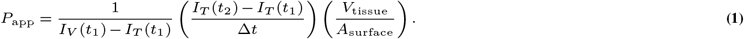

### Cytokine analysis

Supernatants were collected and immediately stored at − 80°C until measurement. For measurement of cytokines (CXCL12, C3, CXCL1, CXCL2, IL6, CCL2, ICAM-1) the customized Invitrogen ProcartaPlex Multiplex Immunoassay (Assay ID: PPX-07-MXEPVUA) was applied and used according to the manufacturer’s instructions. Capture beads were added to a 96-well flat-bottom plate, then washed using a Bioteck 405TS microplate washer. Next, the beads were incubated for 2 hours at room temperature with either diluted supernatants or provided standards. After another wash, detection antibodies were added and incubated for 30 minutes at room temperature, followed by a final wash. The beads were then incubated with Streptavidin-PE for 30 minutes at room temperature before the last washes. Finally, the beads were resuspended in an acquisition buffer and the plate was read on a Bio-Rad Bio-Plex 200 instrument using Bio-Plex Manager Software v. 6.2.

### Dissociation of the Fibrin Gel within the Microfluidic Chip

In order to remove the cells attached to the media channel, the media channels were washed with DPBS (Gibco, − CaCl_2_, − MgCl_2_) supplemented with 2 mM EDTA (Add supplier) by adding 70 µl to one side and 50 µl to the other to induce parallel flow. After volume equilibration, this washing step was repeated. Next, 0.25% trypsin (Gibco) containing DNAase (Roche) was added with parallel flow, as described above. Once the volumes equilibrated, the trypsin-DNAase mixture was added again in the same manner. The chips were then placed in a humidified incubator (37°C, 5% CO_2_) for 4–6 min. Once the cells in the side channel were detached and the vascular network remained intact, the trypsin-DNAase solution was aspirated using a vacuum pump.

After aspiration, the enzyme mix was added with a parallel flow as described previously. The microfluidic chips were subsequently placed in a humidified incubator (37°C, 5% CO_2_). The integrity of the vascular network was monitored under the microscope every two minutes. Upon observing the onset of compromise to its integrity, the enzyme mix was removed, and nattokinase solution was introduced with interstitial pressure into the media ports. Additionally, it was added into the gel inlet and outlet. The microfluidic chips were then incubated in a humidified incubator (37°C, 5% CO_2_) for 25–30 min on the OrganoFlow rocker, oriented perpendicularly to the rocker’s axis. The rocking interval was 2 min at an angle of 15°. The nattokinase solution was pipetted through the inlet/outlet gel port every 10 min, with gel dissociation checked under the microscope after each pipetting step.

Subsequently, the gel was removed from the microfluidic chip with a pipette through the gel inlet/outlet port and transferred to a 15 ml Falcon tube containing 5 ml of pre-warmed enzyme mix. This was incubated on a rocker (1 min interval, angle: 16°) in a humidified incubator (37°C, 5% CO_2_) for 20 min. Every 10 min, the gel was carefully pipetted to mechanically aid dissolution. Once the gel was completely dissolved, cold medium with 10% FBS (Add supplier) was added, and the solution was centrifuged at 4°C (300g, 5 min). After centrifugation, cells were resuspended in cold medium to break up any aggregates, then filtered sequentially through a 70 µm and subsequently a 40 µm filter to obtain a single cell solution.

### Imaging and analysis

Prior to imaging, vascularized microfluidic chips were washed 3× with fresh culture medium to remove flowing PBMC from the vascular network. Endothelial cells (HUVECs) were stably expressing GFP, immune cells were fluorescently labeled with CellTrace™ Far Red (Thermo Fisher), and tumoroids were transfected with an RFP reporter construct, for identification.

Imaging was performed with a Leica SP8 or Stellaris 8 confocal microscope.

Following acquisition, merged files were exported and subsequently processed using Imaris (Imaris x64 9.7.1) for image visualization and analysis.

To assess the expression of the Intercellular Adhesion Molecule (ICAM-1) and vascular cell adhesion molecule 1 (VCAM-1), live cell imaging was performed, following treatment with TNFa (30 ng/mL) for 4 hours. Vascularized microfluidic chips (idenTx9, AIM Biotech) were washed and then infused with ICAM-1 antibody (BioLegend, 1:100) and VCAM.1 antibody (BioLegend 1:100) in media. After a 1-hour incubation, chips were washed with fresh media using a pressure gradient.

### Immunofluorescence Staining

For immunofluorescence staining on vascularized microfluidic chips (idenTx9, AIM Biotech) samples were fixed by perfusion with 4% paraformaldehyde (PFA) in PBS for 15 min at room temperature, followed by two PBS washes. Before staining, slides containing chip sections were prepared by carefully removing the laminate covers using fine tweezers to expose the tissue regions. Samples were immediately covered with 50–100 µL of sterile-filtered blocking buffer (3% FBS, 1% BSA, 0.5% Triton X-100, and 0.5% Tween-20 in PBS) and incubated for at least 2 h at room temperature or overnight a 4°C.

After blocking, primary antibodies were diluted in blocking buffer and incubated overnight at 4°C. The following antibodies were used in this study: CD31 (Dako, M082329-2), EPCAM-1 (Miltenyi Biotech, 130-136-797), Cleaved Caspase 3 (Cell Signaling, 9661), Fibronectin (Sigma, F3648), CollagenV (Abcam, ab321795), CollagenVI (Abcam, ab182744), SMA (Sigma, A2542), Transgelin (Abcam, ab14106), FAP (Invitrogen, BMS168). Chips were sealed to prevent evaporation and incubated overnight at 4°C. On the next day, following three washes with PBS, 20 min each, secondary antibodies (1:250 in blocking buffer) were added and incubated for 2 h at room temperature in the dark. Samples were washed three additional times with PBS-T for 30 min. Slides were mounted using Vectashield Vibrance mounting media and allowed to cure for 1–2 h at room temperature before storage at 4°C until imaging.

### RNA sequencing

Prior to the preparation of single cell RNA sequencing libraries the amount of cells in the single cell suspensions was determined and the cell viability was measured using a Countess II cell counter (Invitrogen/Thermo Fisher Scientific). The cell viability of all samples was high and ranged between 89% and 98% viability.

ScRNAseq libraries were prepared using the Next GEM Single Cell 3 Reagent Kit v3.1 (10X Genomics) according to the manufacturer’s instructions. Viable single cells were loaded onto a 10x Genomics Chromium Next GEM Chip G and processed in a 10x Genomics Chromium controller with a target range of 8.000 to 12.000 cells per sample. Amplification of the cDNA was performed during 11 PCR cycles. The final sequencing libraries were checked for their quality using the High Sensitivity DNA Kit (Cat # 5067-4626 and # 5067-4627) on the Bioanalyzer (Agilent Technologies, Inc.) and quantified using the Qubit 1X dsDNA HS Assay Kit (Cat #Q33230, Invitrogen/Thermo Fisher Scientific). After normalization, pooling and a PhiX spike in of 1% the libraries were paired-end sequenced on the Illumina NovaSeq6000 instrument (Illumina, Inc.) using S2 flow cells (Cat # 20028316, 100 cycles) and a run protocol of 28 cycles for read 1 and 90 cycles for read 2. The libraries were sequenced with an average of 30.000 reads per cell.

The raw sequencing reads were processed by Cell Ranger (v. 6.0.2) with the default parameters and aligned against the human reference genome (GRCh38).

### Data Processing and Analysis

All analyses were performed in R (v4.3) utilizing the Seurat package (v5; Hao et al. 2024) for single-cell RNA-seq data processing. To optimize computational efficiency, BPCell (v0.3; Parks & Greenleaf 2025) was used as the backend for accelerated dimensionality reduction and integration within Seurat. Parallelization was implemented using the future package (Bengtsson 2021).

Raw gene expression count matrices were processed using Seurat. Cells and genes were filtered based on standard quality control metrics in Seurat. Data normalization was performed with the `NormalizeData` function using default parameters. Highly variable genes (*n* = 4000) were identified with `FindVariableFeatures`.

### Dimensionality Reduction and Batch Correction

Gene expression values were scaled (`ScaleData`), followed by principal component analysis (PCA) via `RunPCA`. The number of significant principal components was assessed using elbow plots. To correct for batch effects, we applied fast mutual nearest neighbor (FastMNN; Haghverdi et al. 2018) integration implemented via BPCell, and using the subsequent PC embeddings as the input to `IntegrateLayers`.

### Clustering and Cell Type Assignment

A shared nearest neighbor (SNN) graph was constructed using *k* = 30. Clustering was performed using the Louvain or Leiden (Traag et al. 2018) algorithm across a resolution range (0.1 to 2.0), and resolution of 0.4 was selected for downstream interpretation based on cluster stability and biological interpretability. Major cell types (Fibroblasts, Endothelial, Epithelial, etc.) were assigned based on canonical marker gene expression and expert annotation (Supplementary Table X). Subclustering analyses were subsequently performed within each major cell type to refine cluster identities, guided iteratively by domain expert review (Supplementary Table X).

### Differential Expression and Pathway Enrichment

Cluster and cell type marker genes were identified using Seurat’s `FindAllMarkers` function, which applies the Wilcoxon rank-sum test. Genes were considered markers if they were upregulated in each cluster with minimum fold change of 1.5, and expressed in at least 25% of cells within that cluster.

Gene set enrichment analysis was performed using the fgsea package (v1.26.0; Korotkevich et al. 2019) on differentially expressed gene lists. Pathway enrichment was tested for Gene Ontology (GO) Biological Process, C5, and Hallmark gene sets from MSigDB (Liberzon et al. 2015) using a gene set size range of 10–1000 genes.

---

**Supplementary Video 1. Fibroblasts enable self-assembly of human microvascular networks**. Time-lapse fluorescence microscopy of a 5-day co-culture in a microfluidic device showing human endothelial cells (GFP, green) and fibroblasts (magenta). Endothelial cells sprout, align, and anastomose to form a branching network while fibroblasts migrate, contact endothelial cords, and remodel the surrounding matrix, stabilizing nascent vessels. The movie illustrates sequential steps of network assembly—sprouting, guidance, junction formation, and maturation—culminating in a connected microvascular plexus. Colors correspond to the legend (green: endothelial cells; magenta: fibroblasts). Contrast is adjusted uniformly across frames. This recording is representative of the behavior observed across independent chips.

**Supplementary Video 2** | **Vascularization of patient-derived colorectal cancer organoids on chip**. Time-lapse fluorescence microscopy of a self-assembling vascularized organoid culture showing endothelial network formation and stromal organization around patient-derived CRC organoids. CRC organoids are shown in red, endothelial cells in green, and lung fibroblasts in cyan. The movie illustrates progressive endothelial sprouting and stabilization within the stromal compartment, culminating in a dense vascular meshwork intimately associated with the organoid epithelium (representative field of view).

**Supplementary Video 3. Human PBMC perfusion through the self-assembled vascular network**. Time-lapse of a GFP+ human endothelial network (green) during perfusion of primary human PBMCs (magenta). PBMCs transit through the lumenized microvessels under flow, demonstrating network connectivity and perfusability. Colors: green, endothelial cells; magenta, PBMCs. Movie representative of independent chips/donors.

**Supplementary Video 4. Perfusion through tumor–vascular chips with immune cells**. Time-lapse fluorescence imaging of a co-culture comprising a lumenized human endothelial network (GFP, green), primary human PBMCs (Cell tracker, magenta), and a tumor organoid (RFP, red). PBMCs are perfused through the endothelial microvessels, demonstrating network patency and connectivity as they transit past vessel branches adjacent to the organoid. Colors: green, endothelial cells; magenta, PBMCs; red, tumor organoid. Movie representative of independent chips/donors.

**Supplementary Video 5. Intravascular delivery and circulation of immune cells in vascularized tumor organoids**. Time-lapse fluorescence microscopy showing perfusion of peripheral blood mononuclear cells (PBMCs) through the engineered microvascular network surrounding tumor organoids on chip. Tumor organoids are shown in magenta and endothelial vessels in cyan. PBMC trajectories are rendered in according to their perfusion speed (blue=low, red=high), illustrating intravascular circulation through the vascular bed and dynamic immune-cell transport in the vicinity of the organoid tissue (representative field of view).

**Supplementary Video 6. TNF**α**-activated vasculature displaying leukocyte trafficking behaviors**. Imaging of a human endothelial network on chip after stimulation with TNFα (35 ng mL-1). Endothelial cells are shown in cyan and perfused PBMCs in magenta. An analysis overlay highlights distinct behaviors detected by automated segmentation: stable leukocyte adhesion (purple), transcytosis across the endothelial monolayer (orange), and extravasation into the surrounding matrix (red). The video illustrates the spatial distribution of these events across the network, demonstrating robust inflammatory activation upon TNFα treatment. Colors as indicated.

**Supplementary Video 7. Dextran perfusion of a human microvascular bed for permeability assessment**. Time-lapse of a lumenized endothelial network during introduction of fluorescent Dextran 70 kDa from the inlet (left). The advancing intensity front fills the interconnected capillary plexus, revealing rapid network connectivity and spatially heterogeneous flow paths. The on-screen timer indicates elapsed time from the start of perfusion in minutes. Contrast is applied uniformly across frames; representative of ≥ 3 independent chips.

**Supplementary Video 8. Tumor organoids increase microvascular permeability to 70-kDa dextran**. Time-lapse perfusion of fluorescent Dextran (70 kDa) through human microvessels. Top panel: vascular bed alone shows lumen-confined tracer with minimal leakage. Bottom panel: co-culture with tumor organoids exhibits widespread extravasation and interstitial pooling of the tracer surrounding organoids, indicating elevated endothelial permeability. The on-screen timer denotes elapsed time from the start of perfusion. Acquisition settings and flow conditions are identical between panels; movie representative of independent chips.

**Supplementary Video 9. Quantitative mapping of tumor infiltration and killing on-chip**. Wide-field montage of the tumor–vascular compartment showing a lumenized endothelial network (grey), tumor organoids (cyan), and perfused PBMCs (magenta). The movie overlays an analysis layer that quantifies immune infiltration (extravasated PBMCs, yellow) into and around organoids (tumor-engaged PBMCs, red) and marks caspase-3–positive (orange) tumor cells as apoptotic “killing” events. Per-frame readouts summarize cell counts and infiltration depth and highlight the spatial distribution of apoptosis across the chip. Acquisition and display settings are identical throughout; representative of independent chips/donors.

**Supplementary Video 10. Perfusion of healthy colon organoids**. Wide-field time-lapse during introduction of 70-kDa fluorescent dextran from the inlet (left). On-screen timer denotes elapsed time from the start of perfusion. Acquisition and display settings are identical across frames; representative of independent chips/donors.

**Extended Data Fig. 1.**
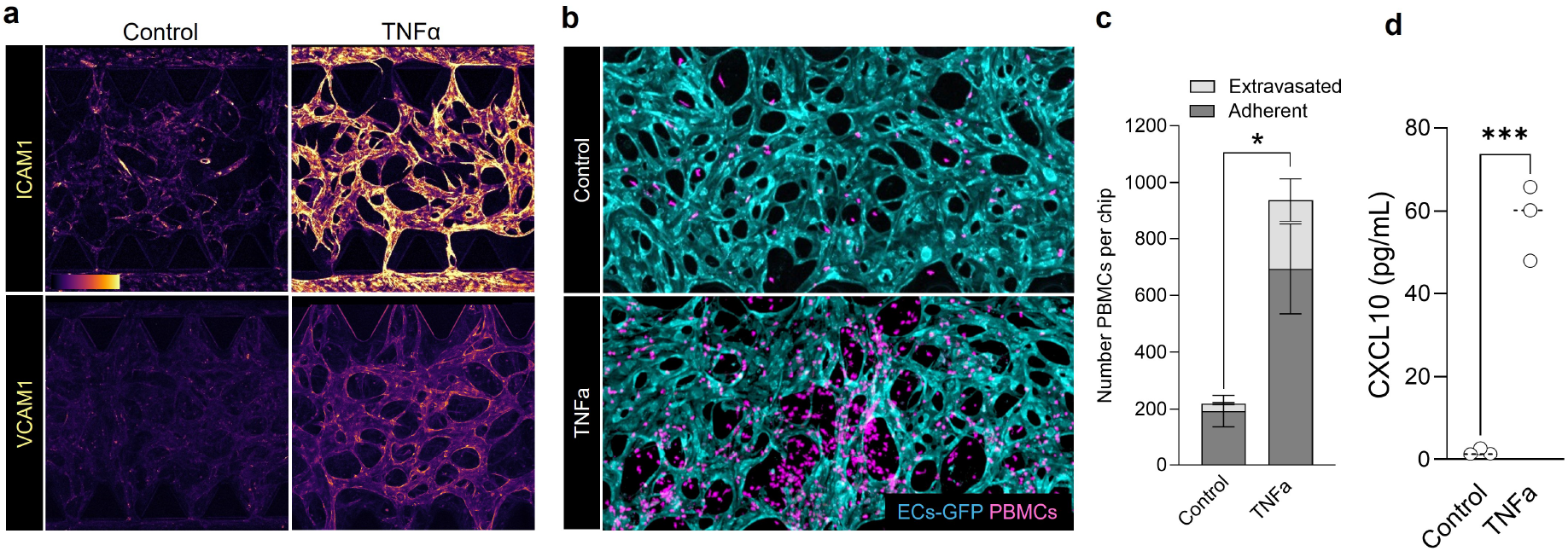
Recapitulating inflammatory response and immune trafficking on-chip. a, Endothelial activation by TNFα induces robust up-regulation of adhesion molecules across the microvascular network, shown as intensity heat maps for ICAM1 and VCAM1 (Control vs TNFα). b, Representative confocal images of the vascularized channel showing endothelial cells (cyan) and PBMCs (magenta/red). TNFα priming increases leukocyte tethering/adhesion and transendothelial migration (white dashed outlines; higher-magnification insets). c, Quantification of leukocyte–vessel interactions per chip, partitioned into adherent (grey) and extravasated (magenta) cells; TNFα markedly elevates both behaviors (bars = mean, error bars = s.d.; points = individual chips). d, Secreted CXCL10 measured from chip effluents rises after TNFα stimulation (points = chips; bars = mean ± s.d.). Data are representative of independent chips per condition.

**Extended Data Fig. 2.**
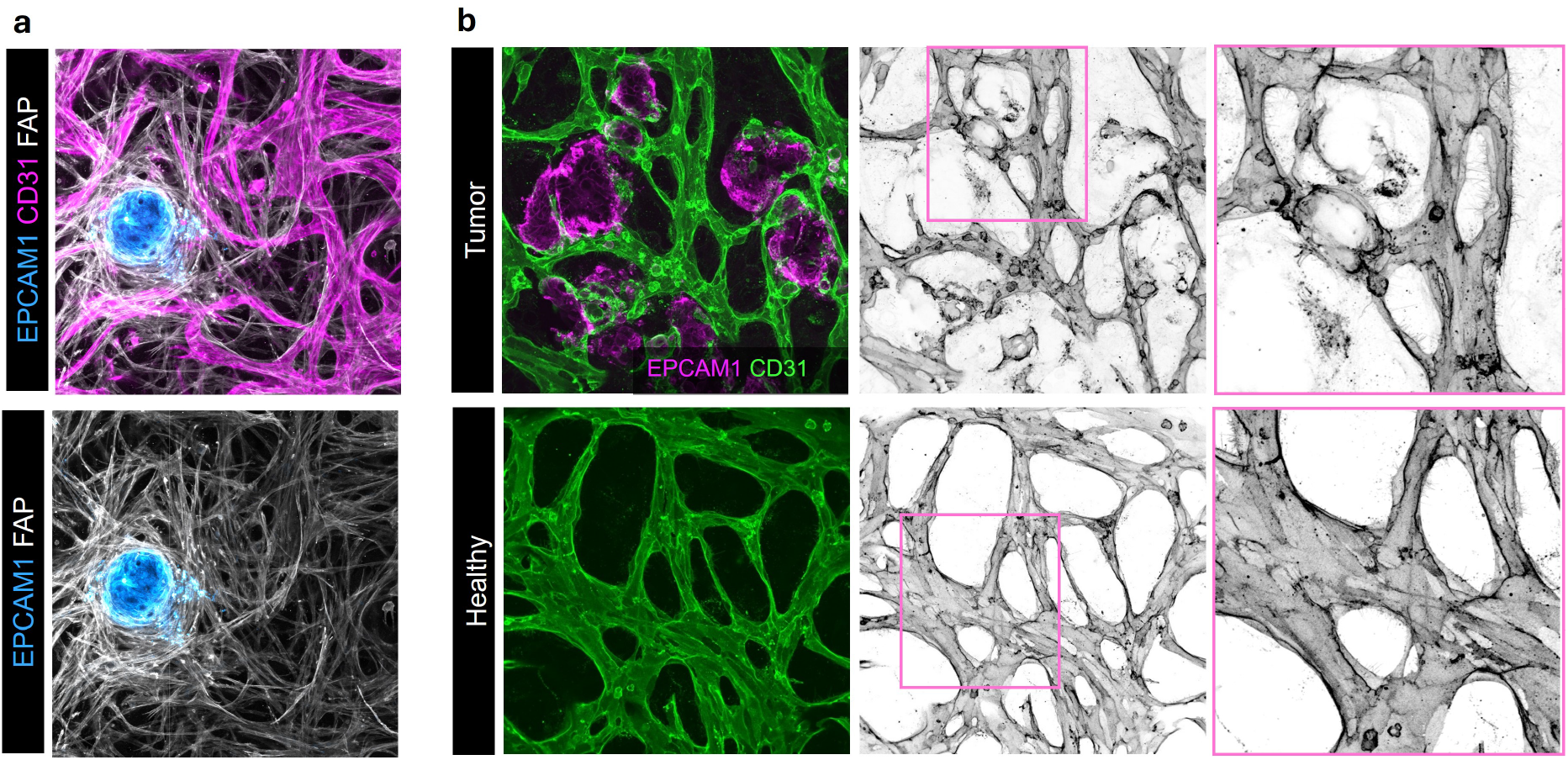
Tumor organoids promote FAP fibroblast accumulation at the tumor–stroma interface and induces disorganized vascular architecture. a, Representative confocal maximum-intensity projection of VascO showing the tumoroid (blue), endothelial cells (ECs; magenta) and FAP immunofluorescence (white/grey). FAP fibroblasts form a peritumoral sheath and align with endothelial cords. b, Same field with the EC channel removed to highlight the peritumoral FAP cuff around the tumoroid (blue). Data are representative of independent chips. b, Representative confocal images of vascularized chips with tumor organoids (top row) or endothelium alone (bottom row). Left panels show EPCAM1 tumoroids (magenta) and CD31 endothelium (green); middle panels display the CD31 channel in grayscale; right panels are magnifications of boxed regions. In the presence of tumoroids, vessels exhibit irregular calibres, tortuous branches, blind ends and exuberant sprouting, whereas endothelium cultured without tumoroids forms a more regular, reticular network. Images are representative of independent chips.

**Extended Data Fig. 3.**
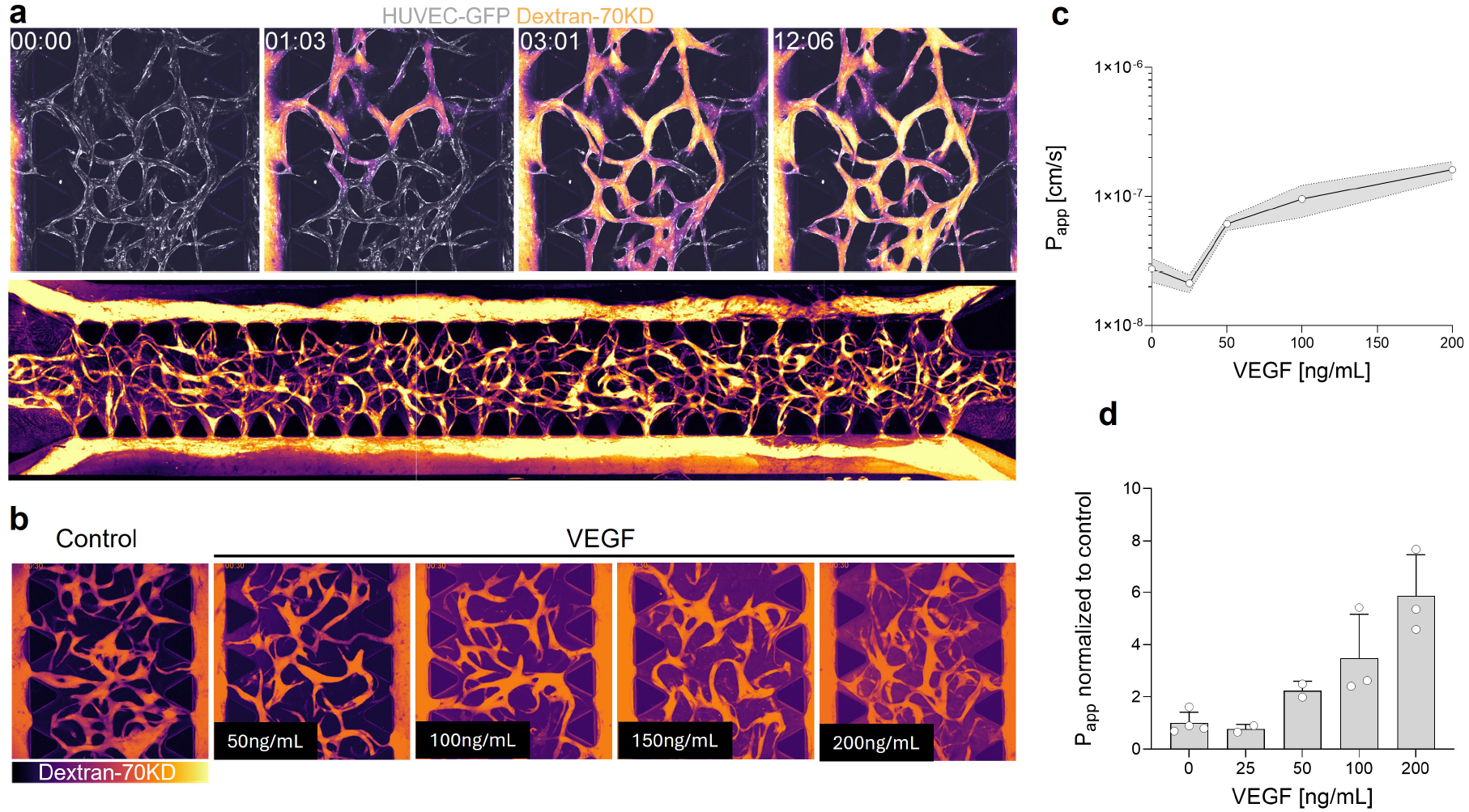
Recapitulation of VEGF-induced vascular barrier disruption. a, Time-lapse montage of fluorescent 70-kDa dextran perfusion through the endothelial network (pseudocolor; yellow/white = higher intensity) with frames at 0:00, 1:03, 3:01 and 12:06 min, followed by a wide-field view showing complete chip perfusion. b, Representative endpoint images after exposure to the indicated VEGF concentrations (0–200 ng mL-1), illustrating dose-dependent barrier breakdown. c, Apparent permeability (Papp) estimated from time-lapse sequences increases with VEGF in a dose-responsive manner (line = mean; shaded band = variability across fields/chips). d, Papp normalized to matched controls for each experiment, plotted against VEGF dose (points = individual fields/chips; horizontal bars = mean ± s.d.).

**Extended Data Fig. 4.**
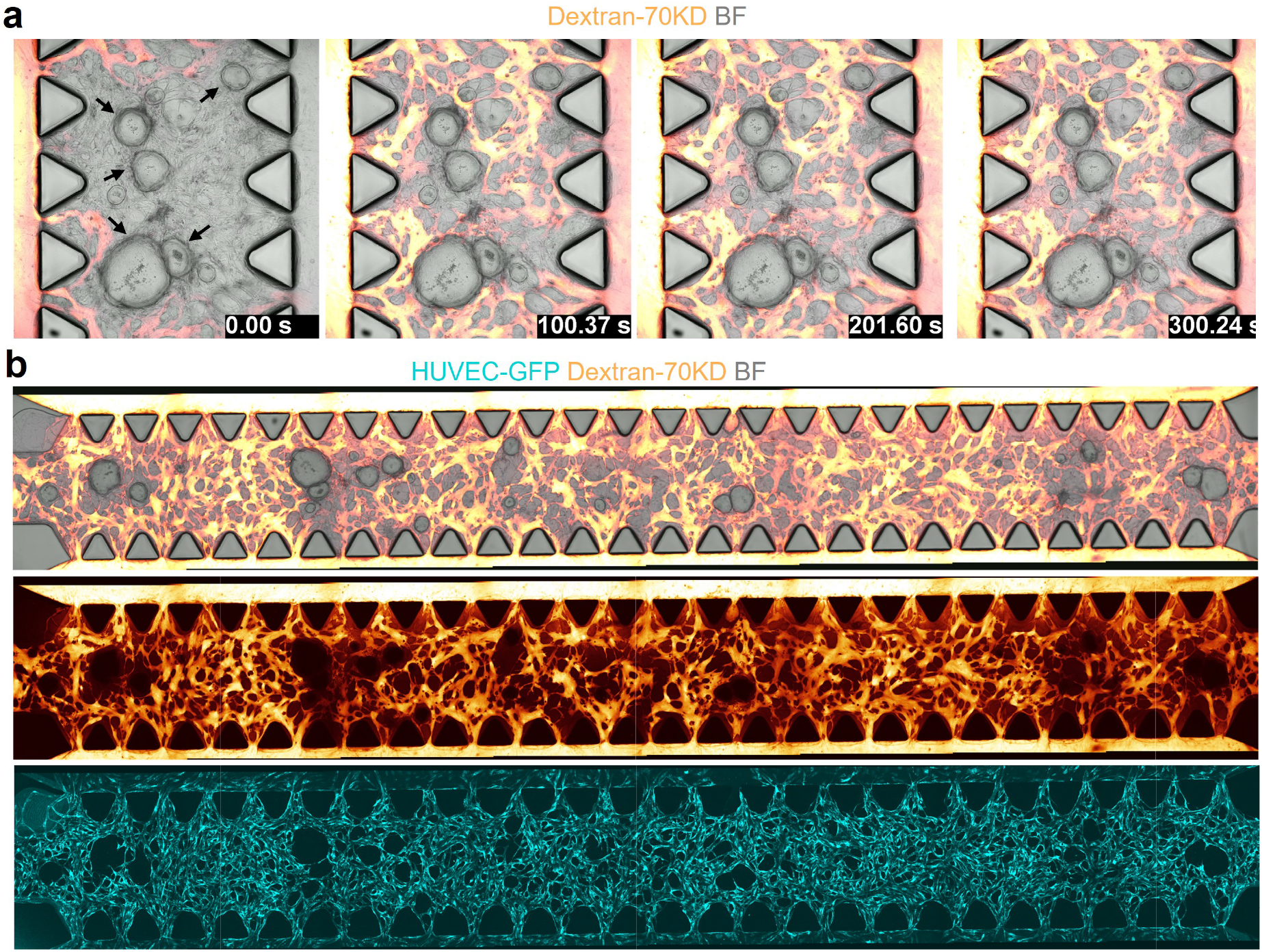
Perfusion of healthy colon organoids. a, Time-lapse montage of a microvascular network co-cultured with healthy colon organoids during perfusion of fluorescent 70-kDa dextran (pseudocolor; yellow/white = higher intensity) at 0, 100, 200 and 300 s,showing rapid intraluminal filling adjacent to organoids. b, Wide-field views of the perfused channel along the chip with (top) and without (bottom) organoids illustrate uniform tracer distribution throughout the network. Images are representative of independent chips.

